# Mu Transposition in the Absence of the Target-capture Protein MuB Reveals New Roles of MuB in Target Immunity and Target Selection, and Redraws the Boundaries of the Insular Ter Region of *E. coli*

**DOI:** 10.1101/2020.04.24.060434

**Authors:** David M. Walker, Rasika M. Harshey

## Abstract

The target capture protein MuB is responsible for the high efficiency of phage Mu transposition within the *E. coli* genome. However, some targets are off-limits, such as regions immediately outside the Mu ends (*cis*-immunity) as well as the entire ∼37 kb genome of Mu (Mu genome immunity). Paradoxically, MuB is responsible for *cis*-immunity and is also implicated in Mu genome immunity, but via different mechanisms. In this study, we tracked Mu transposition from six different starting locations on the *E. coli* genome, in the presence and absence of MuB. The data reveal that Mu’s ability to sample the entire genome during a single hop in a clonal population is independent of MuB, and that MuB is responsible for *cis*-immunity, plays a lesser role in Mu genome immunity, and facilitates insertions into transcriptionally active regions. Unexpectedly, transposition patterns in the absence of MuB have helped extend the boundaries of the insular Ter segment of the *E. coli* genome.

## Background

Phage Mu uses transposition to amplify its genome ∼100-fold during its lytic cycle in *E. coli*, making it the most efficient transposable element (TE) described to date [1-3] (Fig. 1A). Mu transposes by a nick-join pathway, where assembly on Mu ends of a six-subunit MuA transposase complex (transpososome) is followed by introduction of nicks at both ends; the liberated 3′-OH groups at each end then directly attack phosphodiester bonds spaced 5 bp apart in the target DNA, covalently joining Mu ends to the target [4]. The resulting branched Mu-target joint is resolved by replication, duplicating the Mu genome after every transposition [5]. At the end of the lytic cycle, Mu copies are excised for packaging by a headful mechanism that cuts and packages host DNA on either side of Mu [1, 6]. The latter finding has been exploited to examine target site preference *in vivo* by sequencing the flanking host DNA packaged in Mu virions [7, 8].

**Figure 1.**
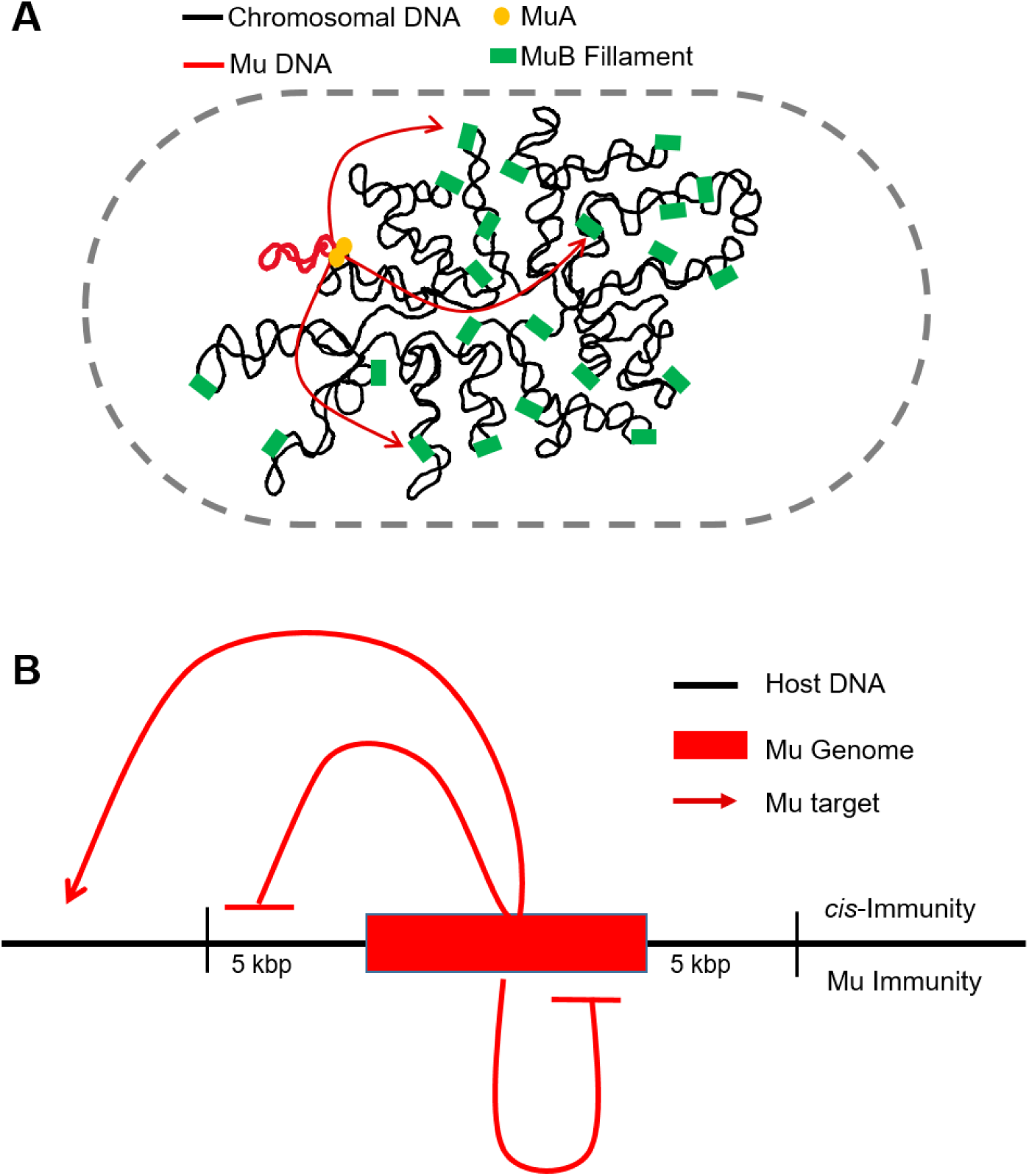
Mu transposition and target immunity. **A**. The transposase MuA pairs Mu ends and introduces single-strand nicks, joining these to MuB-captured target DNA. MuB binds DNA non-specifically, polymerizing in short filaments, and increases the catalytic efficiency of target capture. **B**. Both *cis-*immunity and Mu genome immunity operate by two distinct mechanisms to prevent Mu insertion. *Cis-*immunity is characterized by the lack of insertions outside Mu ends, typically within 5 kb. Mu genome immunity is characterized by absence of insertion anywhere within the ∼37 kb Mu genome.

The B protein of Mu (MuB), a non-specific DNA-binding protein and AAA+ ATPase, is essential for the efficient capture and delivery of the target to the transpososome via MuB-MuA interaction; MuB also plays critical roles at all stages of transposition by allosterically activating MuA (see [3, 9]). MuB forms ATP-dependent helical filaments, with or without DNA [10-12]. Because of a mismatch between the helical parameters of the MuB filament and that of the bound DNA, it has been proposed that the DNA at the boundary of the MuB filament deforms, creating a DNA bend favored by MuA as a target [11, 13, 14]. While most TEs display some degree of target selectivity [15], Mu is perhaps the most indiscriminate, with a fairly degenerate 5bp target recognition consensus [7, 8, 16, 17]. Even though MuB facilitates target selection, recognition of the pentameric target site is a property of MuA, and is independent of MuB.

Several bacterial TEs, including members of the Tn*3* family, Tn*7*, and bacteriophage Mu, display transposition immunity [15, 18-22]. These elements avoid insertion into plasmid DNA molecules that already contain a copy of the transposon (a phenomenon called *cis*-immunity), and it is thought that this form of self-recognition must also provide protection against self-integration (TE genome immunity) (Fig. 1B). While *cis*-immunity *in vitro* extends over the entire plasmid harboring the TE, it does not provide protection to the entire bacterial genome on which the TE is resident, but can extend over large distances from the chromosomal site where it is located. *In vitro* studies with mini-Mu donor plasmids provided the first molecular insights into the *cis*-immunity phenomenon [9, 23]. Ensemble and single-molecule experiments showed that MuB bound to DNA dissociates upon interaction in *cis* with MuA bound to the Mu ends, resulting in depletion of MuB near the vicinity of Mu ends, making the depleted region a poor target for new insertions [24, 25]. It was assumed that this mechanism also protects DNA inside Mu ends. *Cis*-immunity has been observed *in vivo*, appearing strongest around 5 kb outside the Mu ends, and decaying gradually between 5 and 25 kb [26, 27].

The proposition that *cis*-immunity also prevents self-integration is a reasonable one for TEs whose size is smaller than the range over which this immunity extends. For Mu, *cis*-immunity has been tested over a 2-3 kb range *in vitro* using mini-Mu plasmids, and found to be strongest around 5 kb *in vivo* as stated above. The range of immunity seen *in vivo* would not be expected to effectively protect the 37-kb Mu genome by the *cis*-immunity mechanism, as was indeed demonstrated to be the case [27]. Therefore, a distinct ‘Mu genome immunity’ mechanism was proposed to explain the lack of self-integration. Unlike the *cis*-immunity mechanism, which requires removal of MuB from DNA adjacent to Mu ends, MuB was observed to bind strongly within the Mu genome during the lytic cycle, suggesting that the mechanism of Mu genome-immunity must be different from that of *cis*-immunity [27]. ChIP experiments revealed sharply different patterns of MuB binding inside and outside Mu, leading to a proposal that the Mu genome is segregated into an independent chromosomal domain *in vivo* [27]; this proposal was confirmed by Cre-*loxP* recombination and 3C experiments for Mu prophages at two different *E. coli* chromosomal locations [28]. A model for how the formation of an independent “Mu domain” might nucleate polymerization of MuB on the genome, forming a barrier against self-integration, was proposed [27].

The present study investigates the role of MuB in the three diverse functions discussed above - target capture, *cis-*immunity, and Mu genome immunity *in vivo*. Through comparison of insertion patterns of wild-type (WT) Mu and MuΔB prophages placed at six different locations around the *E. coli* genome, we show that *cis*-immunity depends on MuB, while Mu genome immunity is only slightly breached in its absence. The data also reveal a previously unappreciated role for MuB in facilitating Mu insertion into transcriptionally active regions, as well as several interesting and hitherto unknown aspects of Mu target choice *in vivo*. An unanticipated outcome of this study is the finding that the Ter segment of the *E. coli* genome, which is more isolated from the rest of the genome, is larger than previously estimated.

## Results and Discussion

### Mu samples the entire *E. coli* genome even in the absence of MuB, helping define new boundaries for the Ter region

We recently exploited the DNA-DNA contact mechanism of phage Mu transposition to directly measure *in vivo* interactions between genomic loci in *E. coli* [29]. Thirty-five independent Mu prophages located throughout the genome were allowed to go through one round of transposition. The data showed that in a clonal population, Mu is able to access the entirety of the genome with roughly equal probability regardless of its starting genome location, suggesting widespread contacts between all regions of the chromosome. The data led us to conclude that the chromosome is well-mixed and shows a ‘small world’ behavior, where any particular locus is roughly equally likely to be in contact with any other locus. The exception was the Ter region, reported by Mu as being less well-mixed than the rest of the genome.

While MuB is essential for target capture *in vitro* [4], transposition is still detectable *in vivo* in the absence of MuB at an efficiency nearly two orders of magnitude lower than WT [30]. To examine how MuB influences the target selection *in vivo*, we monitored insertion patterns of a subset (six) of the Mu prophages used in the original study [29] (Fig. 2A), after a single round of transposition, in the presence and absence of MuB (WT vs ΔMuB) (Fig. 2B). For analysis, the genome was partitioned into 200 equally sized bins (each bin ∼23.2 kb) (Fig. 2A). To generate sufficient insertion resolution, transpositions were analyzed using a target enrichment protocol [29] and deep sequencing of 10 million reads or more. Due to lower transposition frequencies of ΔMuB prophages, these were sampled ∼50% more with a 15 million read depth. The data plotted in Figure 2B show similar insertion profiles for both WT and ΔMuB throughout the genome after normalizing to the read depth for both prophages. Thus, like WT, the ΔMuB prophages transpose to every bin of the genome in a clonal population, allowing us to conclude that the ability of Mu to sample the entirety of the genome in one transposition event is independent of MuB.

**Figure 2.**
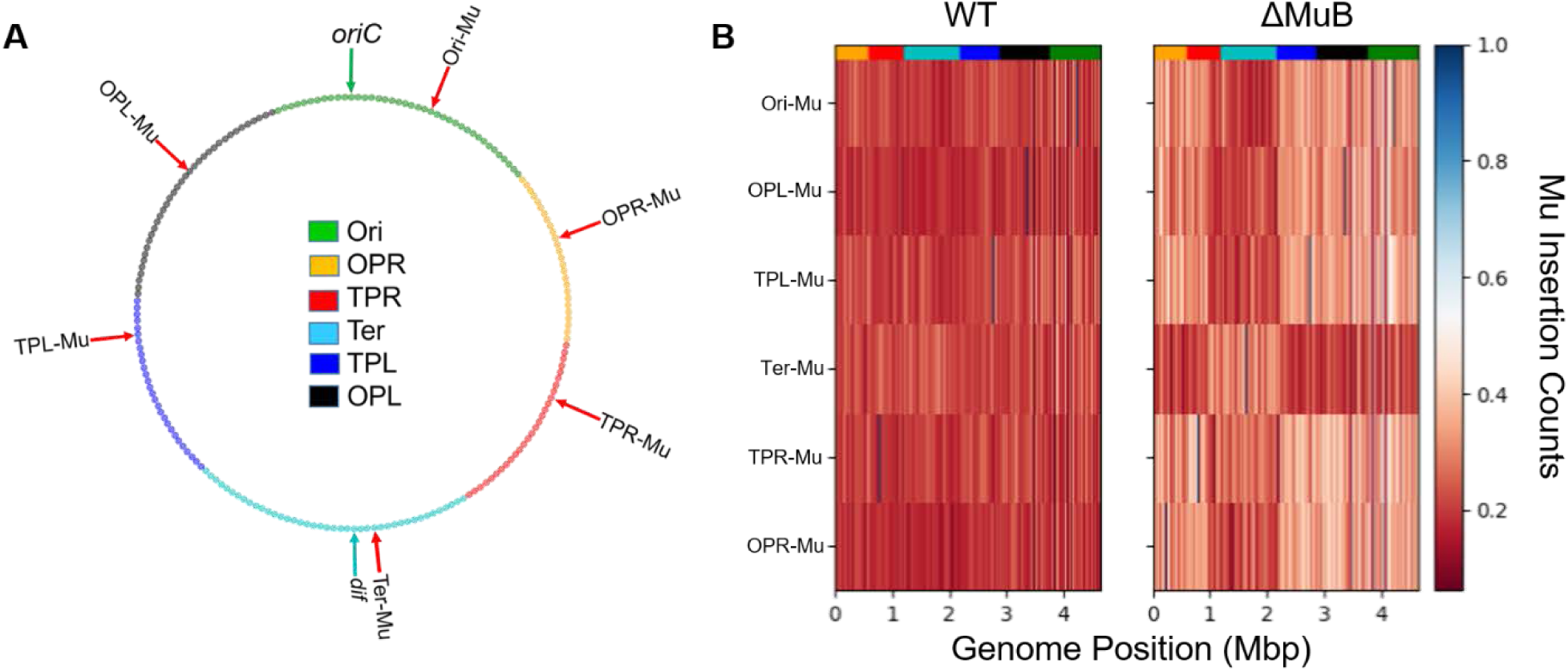
Mu samples the entire genome regardless of the presence of MuB. **A**. The six starting prophage locations on the *E. coli* genome monitored in this study are indicated by red arrows (see Table 1 for their exact locations). These locations were chosen because they are spread throughout the chromosome, and therefore ideally suited for sampling features across the genome. *oriC* in the Ori region is the site where bi-directional replication begins (green arrow), terminating at the *dif* site, exactly opposite to *oriC* within the Ter region (cyan arrow). OPL, Ori proximal left; OPR, Ori proximal right; TPL, Ter proximal left; TPR, Ter proximal right. The boundaries of the various colored regions are taken from [1]. **B**. The genome was partitioned into 200 equally sized bins (A), and the normalized number of unique insertions into each bin for each prophage was computed, as displayed by the color bar. The highest number of unique insertions for any non-starting bin was ∼8000 insertions corresponding to just under 1.0. Each starting bin position can be identified by its high number of counts (deep blue bins); the large number of reads associated with the starting position information were retained to aid in identifying this initial position shown on the map in **B**. The multi-color strip on top of each panel corresponds to chromosomal the regions shown in A. The Ter region (cyan) as explored by the Ter-ΔMuB prophage is 217 kbp larger than earlier estimates [1]. This is recognizable as a square block of lighter red insertions in the Ter-ΔMuB prophage, which lines up with identical blocks of darker red insertions in the other five ΔMuB prophages.

The color-coded map of the *E. coli* genome shown in Figure 2A depicts the length and boundaries of chromosomal regions deduced by prior methodologies to be either unreactive or partially reactive with the other regions [31]. With the exception of Ter, the Mu methodology failed to detect all such boundaries [29]. The Ter region has unique properties shaped by the activity of MatP [32] and the condensin MukBEF [33, 34], and has been shown by several methodologies to be more isolated from the rest of the chromosome [29, 35, 36]. Comparison of WT vs ΔMuB insertion patterns supported this conclusion while revealing more details. For example, the ΔMuB prophage located in Ter (Ter-Mu) had >40% of its total insertions occur within the Ter region, which only comprises ∼20% of the genome (light red profile). While Ter-ΔMuB prophage sampled the DNA around its starting location more efficiently than it did the rest of the genome, the ΔMuB prophages at the five other locations showed a converse pattern in that they could not access Ter as easily (dark red profile). The latter prophages had <15% of their total insertions within Ter. Comparison of both the outgoing and incoming ΔMuB profiles all lined up precisely, giving us a clearer view of the boundaries flanking Ter. According to Valens et al. [31], the Ter region extends from nucleotide position 1128 kb (26’ on the genetic map) to 2038 kb (47’). According to the transposition patterns of ΔMuB prophages, the Ter region extends from nucleotide position 911 kb (21’) to 2200 kb (47’), expanding the left boundary by more than 217 kb (Fig. 2B). We note that ΔMuB prophages did not reveal other boundaries (as demarcated by the colored segments in Fig. 2A) proposed by prior methodologies [31].

Why does such a defined Ter segment emerge only in MuB-deficient prophages? Given that the ΔMuB prophage in Ter had no trouble sampling within Ter, but that the other ΔMuB prophages did have difficulty inserting here, we suggest that the answer lies in the existence of some special feature at the Ter boundaries that isolates Ter. MatP, which binds to specific matS sequences distributed within Ter [32], has been shown to functionally exclude the SMC/condensin complex MukBEF from Ter [33]. Fluorescence experiments have shown that the extent to which MatP organizes Ter and excludes MukBEF ranges from 852 kb to 2268 kb, which is much more in line with our estimates of Ter in the ΔMuB prophages [34]. To our knowledge, MatP is not itself enriched at the Ter boundaries [34]. Perhaps, as an SMC complex, with assistance from other proteins, MukBEF tethers the two chromosomal arms at the Ter boundary, preventing Ter from mixing with the rest of the genome. Given that WT prophages are not as impaired as ΔMuB prophages in sampling Ter, it follows that MuB must weaken the Ter boundary conditions. The property of MuB to nucleate as helical filaments on DNA [11], may be responsible for displacing the boundary-guards. These results imply that the Ter segment is even less well-mixed than determined in the study utilizing WT Mu [29].

### MuB facilitates transposition into highly transcribed regions

Two prior microarray data sets have shown a negative correlation between transcription and Mu transposition [37, 38], although one these studies found several exceptions to this rule, and suggested that some other cellular feature controls these insertion events [38]. We examined this issue using our higher-resolution data set. Fig. 3 compiles a list of 28 genes, most of which are highly transcribed, except for the *lac* operon which is expected to be only partially transcribed under our growth conditions. The figure also includes the flagellar master regulator gene *flhD* which has multiple promoters [39], and *dnaJ* which has no promoter and is exclusively co-transcribed with *dnaK* [40]. For all genes, the earliest identified nucleotide in the coding sequence (CDS) from the annotated genome from genebank (genid: 545778205) is defined as the +1 nucleotide (nt) of the CDS.

**Figure 3:**
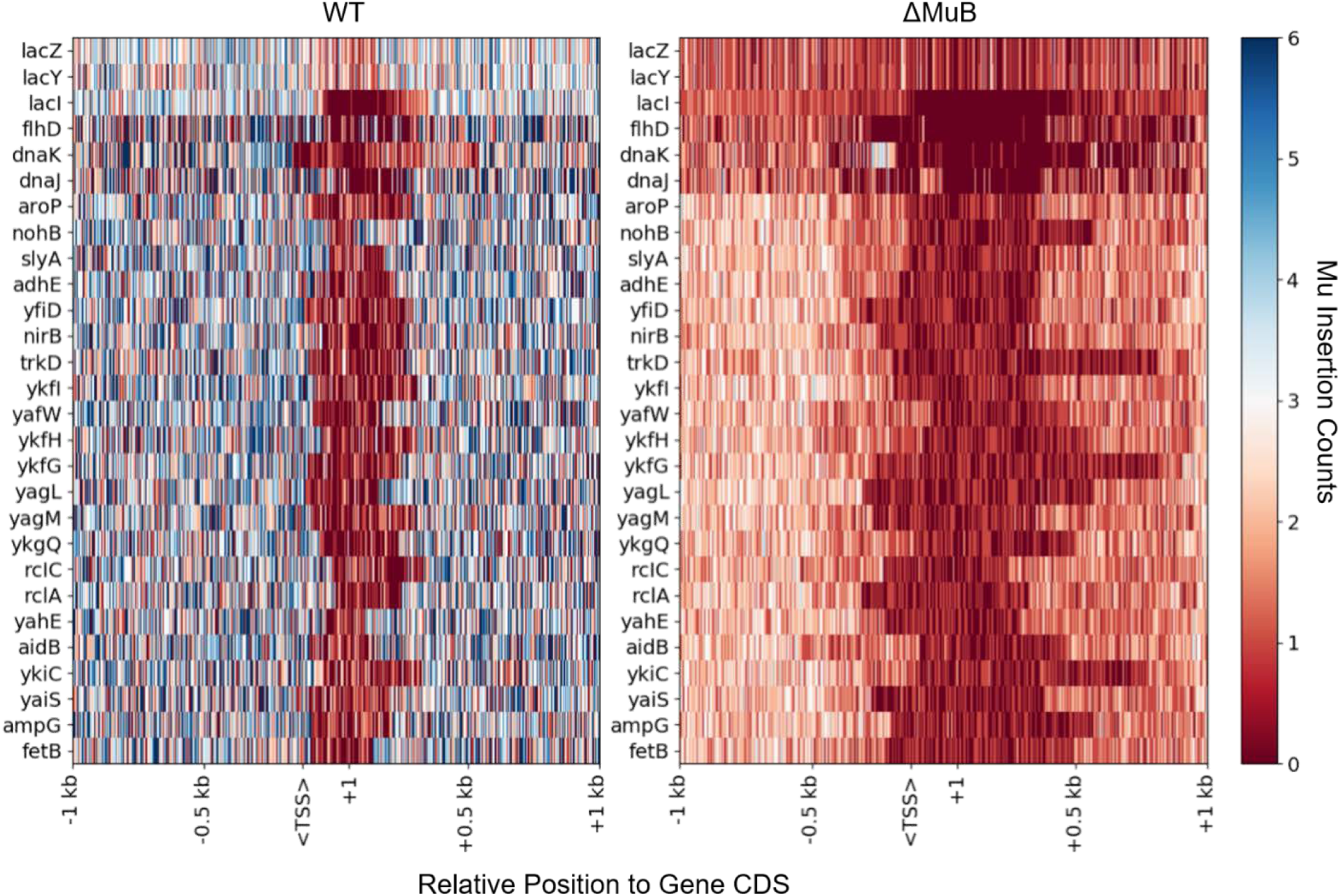
MuB is responsible for capturing target sites near highly transcribed/translated genes. Twenty-three highly transcribed genes, plus the *lac* operon, *flhD* and *dnaK-dnaJ*, were selected for comparison between WT and ΔMuB insertion patterns. For WT transposition, the EST counts (early stage transposition; 15 min induction of transposition) were pooled together from all six prophage locations with an average of 5 million reads per prophage. ΔMuB experiments pooled all six prophages with an average of 20 million reads per prophage. Each gene is oriented to where the +1 nt of coding sequence of the gene starts at the tick mark labeled +1, and downstream nucleotides follow to the right. Upstream nucleotides are marked by negatively labeled tick marks. The expected transcription start site labeled <TSS> is 125 nt away from the +1 site.

WT Mu had significant difficulty inserting near the +1 nt of all active genes, in a region that extends up to 50-200 bp upstream, typically including promoter regions (TSS) [41], and 50-300 bp downstream. However, the transposition difficulty was exacerbated in ΔMuB prophages, which showed an increase in an exclusion zone starting near the TSS for transcriptionally active genes and to a lesser extent for the comparatively less transcriptionally active *lacZY*. Interestingly, two different WT Mu insertion patterns were observed within the *lac* operon, whose *lacZ* and *lacY* genes are repressed by the activity of the *lacI* repressor, which is expected to be transcribed [42]. The number of Mu insertions in *lacI* were roughly half those in *lacZY*, with a strong suppression of insertions around the TSS and +1 nt region of *lacI* for WT. This observation is in agreement with the previous findings of a negative correlation between transcription and transposition.

Of six potential promoters in the *flhDC* operon that control flagellar gene transcription in *Salmonella*, only two (P1 and P5) were seen to be functional [39]. These two sites are each 200-300 bps upstream of the +1 nt [39]. On the other hand, the specific transcriptional start site for *dnaJ* is 2 kb away, as *dnaJ* is always co-transcribed with *dnaK*, with a small 370 nt RNA candidate *tpke11* between the two genes [40, 43]. WT prophages show a near uniform sampling across *flhD*, with reduced insertion around the TSS, while ΔMuB prophages show in addition a secondary exclusion zone upstream from the +1 that encompasses both P1 and P5 promoter regions. Even though TSS is absent in *dnaJ*, WT Mu shows an insertion exclusion zone around +1 of this gene. ΔMuB prophages show an exclusion zone upstream of *dnaJ* not seen in WT, around the position of *tpke11*, while revealing an unusually permissive region upstream of *dnaK*. The latter permissive region in both WT and ΔMuB corresponds to the 377 bp intergenic region between *yaaI* and the *dnaKJ* operon promoter. While this set insertion patterns overall is consistent with the negative correlation between transcription and transposition, particularly around the TSS and +1 for WT, the insertion patterns in *dnaJ* reveal that the +1 region presents a transposition barrier independent of the promoter region, and is likely reflective of the translation activity of the mRNA near this genomic site given that transcription and translation are coupled in bacteria.

To examine Mu insertion patterns in genes that are transcribed but not translated, we looked at both ribosomal RNA operons and tRNA genes. *E. coli* has 7 ribosomal RNA operons that are highly transcribed [44]. We observed a large variation of insertion profiles in these regions (Fig. S1). For example, the insertion frequency of WT Mu is highest in *rrnA*, uniform across the entire operon, and independent of MuB. *rrnE* and *rrnH* receive more insertions in the 23S compared to the 16S region, and are responsive to MuB. *rrnG* shows a large increase in sampling only at the 5’ end of the 16S region (note that *rrnG* is on the negative strand). There seems to be an equal level of Mu insertion between *rrnB, rrnC*, and *rrnD*. If transcriptional status determines Mu insertion efficiency as concluded from the data in Figure 3, then the insertion patterns observed in the *rrn* operons should reflect this as well. Accordingly, *rrnA* is the least transcriptionally active. While early experiments showed little difference in expression levels between the operons in minimal media (*rrnA* actually was reported to have marginally higher expression levels [44]), more recent experiments reporting promoter activity for the *rrn* operons as measured by binding of Fis, a regulator of *rrn* transcription [45], have determined that *rrnE* has the highest level of activity in minimal media with *rrnA* having relatively low levels of promotion [46]. Our results are more in line with the newer data, in that Mu activity is highest within *rrnA*, and lowest near the promoter region of *rrnE* (Fig. S1). Regardless of the *rrn* operon, there seems to be a small window between the 16S and 23S subunits in each operon that is marked by an increase in insertion frequency. This window contains non-coding sequence as well various tRNA sequences. The latter are highly undersampled by Mu insertions even when they occur elsewhere in the chromosome as discussed below.

Mu insertion patterns into 86 tRNA genes scattered throughout the *E. coli* genome [47], is shown in Figure S2. Mu shows an interesting selectivity for inserting into 30 of these genes, avoiding the region that would ultimately be the mature tRNA sequence, as exemplified by the large hole or gap with no insertions seen in the bottom half of the WT Mu panel. Note that Mu is more actively inserting into the genomic regions associated with the 5’ leader and 3’ tailing sequences of pre-tRNA. This would suggest that there is some genomic feature (fold, DNA-binding protein) that is ultimately protecting this region of DNA from Mu insertion. ΔMuB prophages incidentally were less likely to insert into the entire pre-tRNA sequence, suggesting that the transcriptome machinery provided a much higher barrier of access to the ΔMuB prophages over the WT prophages. Using genome-wide transcription propensity data [48], we were able to compare the levels of transcription for each of the tRNA sequences along with the likelihood that Mu (WT and ΔMuB) would transpose within them. Although the transcriptional information was quantitatively sparse amongst most of the tRNA genes, the accessibility of insertion into 36 tRNAs that are the lowest transcriptionally active genes, and exclusion of insertion into the highest transcriptionally active *ileY* and *selC* (marked with red asterisks), is unmistakable. In these two regions, there are no insertions in the entire pre-tRNA CDS in both WT and ΔMuB.

We conclude that the level of availability of a target for Mu insertion is highly correlated with its transcriptional activity, enhanced in the presence of MuB and suppressed in its absence. The particular difficulty of WT Mu in inserting around the TSS could be a combination of DNA strand in the open complex at this site, occupancy by RNA polymerase, or because promoter regions are A/T rich; MuB is reported to exhibit a tendency to form larger filaments on A/T-rich DNA [10, 49]. MuB binding around promoter regions may block insertion of WT Mu there, as Mu transposition has been observed at the junction of A/T and non-A/T DNA *in vitro* [50], and near the vicinity rather than within, MuB-bound regions *in vivo* [38]. For translated genes, the evidence points to a relationship between transcriptional as well as subsequent translational activity of the mRNA in blocking Mu transposition, as demonstrated by insertion patterns around the +1 position of *dnaJ*. In the case of the transcriptionally, and therefore translationally inactive *lacZY* genes we see that there is no barrier to insertion at the +1 nt site, reinforcing this conclusion. As speculated above for the role of MuB in weakening the Ter boundary, we suggest that the filament-forming property of MuB may dislodge transcribing RNA polymerase and ribosomes from transcriptionally active DNA. The most undersampled regions on the genome are coding regions of tRNA, even though Mu is able to sample the leader sequences of the pre-tRNA coding regions, suggesting that some feature of these regions other than transcription protects them from Mu insertion.

### Target consensus *in vivo*

The 5-bp target recognition site for Mu transposition was derived from *in vitro* experiments to be 5′-CYSRG, and observed to be independent of MuB [16, 17]. In the Mu transpososome crystal structure, a hairpin bend in the target was observed, with the transpososome contacting a 20-25 target segment [13]. Preference for a bent target conformation is supported by other *in vitro* experiments [14, 51]. Analysis of target sequences *in vitro* detected symmetrical base patterns spanning a ∼23 to 24-bp region around the target recognition pentamer, indicative not of an extended sequence preference, but possibly of a structural preference that might facilitate target deformation [17].

*In vivo*, a preference for 5’-CGG as the central triplet was derived from cloning 100 Mu-host junctions from packaged phage particles [8]. To re-examine target preference using our current data set, we pooled first-hop insertion data totaling over 120 million targeted Mu reads for both the WT and ΔMuB constructs. We observed that in the genome, sequences with the triple-’G’ consensus and their reverse compliment were 3-4 times more abundant than the 5’-CYSRG-3’ sequences, explaining the preference for 5’-CGG in the earlier study (Fig. S3A). Sequencing data suggest that there is a 7-fold preference for the 8 possible 5’-CYSRG-3’ consensus sequences over the other 1016 remaining pentamer sequences (Fig. S3B).

### MuB is responsible for *cis*-immunity

The *cis-*immunity phenomenon has been studied *in vitro* exclusively by the Mizuuchi group, from ensemble experiments with mini-Mu plasmids to single molecule experiments with tethered λ DNA [9, 23]. A diffusion ratchet model, in which MuA-MuB interactions form progressively larger DNA loops, was proposed to explain the clearing of MuB near the vicinity of Mu ends, with eventual insertion of Mu at sites distant from the ends [24, 25].

We graphed Mu insertions flanking the ends of each starting position, by pooling information from all six prophages during the first round of transposition, as was done for all prior experiments, but we refer to here as early stage transposition (EST), to distinguish them from late stage transposition (LST) where data were collected after multiple rounds of transposition. For the LST condition, we let the experiment run for 2 hours, which allowed WT to complete its lytic cycle (in ∼50 min) and ΔMuB prophages to accumulate 5 to 10 copies of Mu on average per cell as predicted by genome abundance, assuming an even distribution of Mu copy number among the population. All six prophage strains were used for EST experiments, and one WT plus all six ΔMuB prophages for LST experiments.

During EST, WT Mu (bottom row of all plots) does not transpose within 1.5 kb outside each of the starting Mu positions, consistent with the *cis*-immunity phenomenon (Fig. 4A). That the absence of transposition in this region is not due to an intrinsic resistance to insertion within this DNA, is seen from the pooled profiles of the other prophages for the same region (WT pool). Figure 4B examines this pattern in greater detail. There is a slow increase in the number of insertions from about 2 kb outside both Mu ends, exhibiting a sharp increase around 5 kb, and reaching the average number of insertions for bulk DNA at around 10 kb. This pattern was symmetrical for individual ends (Fig. S4). LST samples for WT OPL-Mu show that *cis*-immunity remains intact through multiple rounds of transposition (Fig. 4C). We observed that the insertion patterns outside Mu ends have a sigmoidal nature in both samples, suggesting that *cis*-immunity is not entirely dependent on linear genomic distance. The previously described ratchet-model suggests that intrinsically clustered MuA would hydrolyze proximal MuB-ATP during dynamic loop formation due to Brownian motion [25]. We propose that the sigmoidal pattern of *cis*-immunity arises from three-dimensional features of the genome, and that dynamic loop formation is the necessary factor in creating the conditions that generate such a pattern.

**Figure 4.**
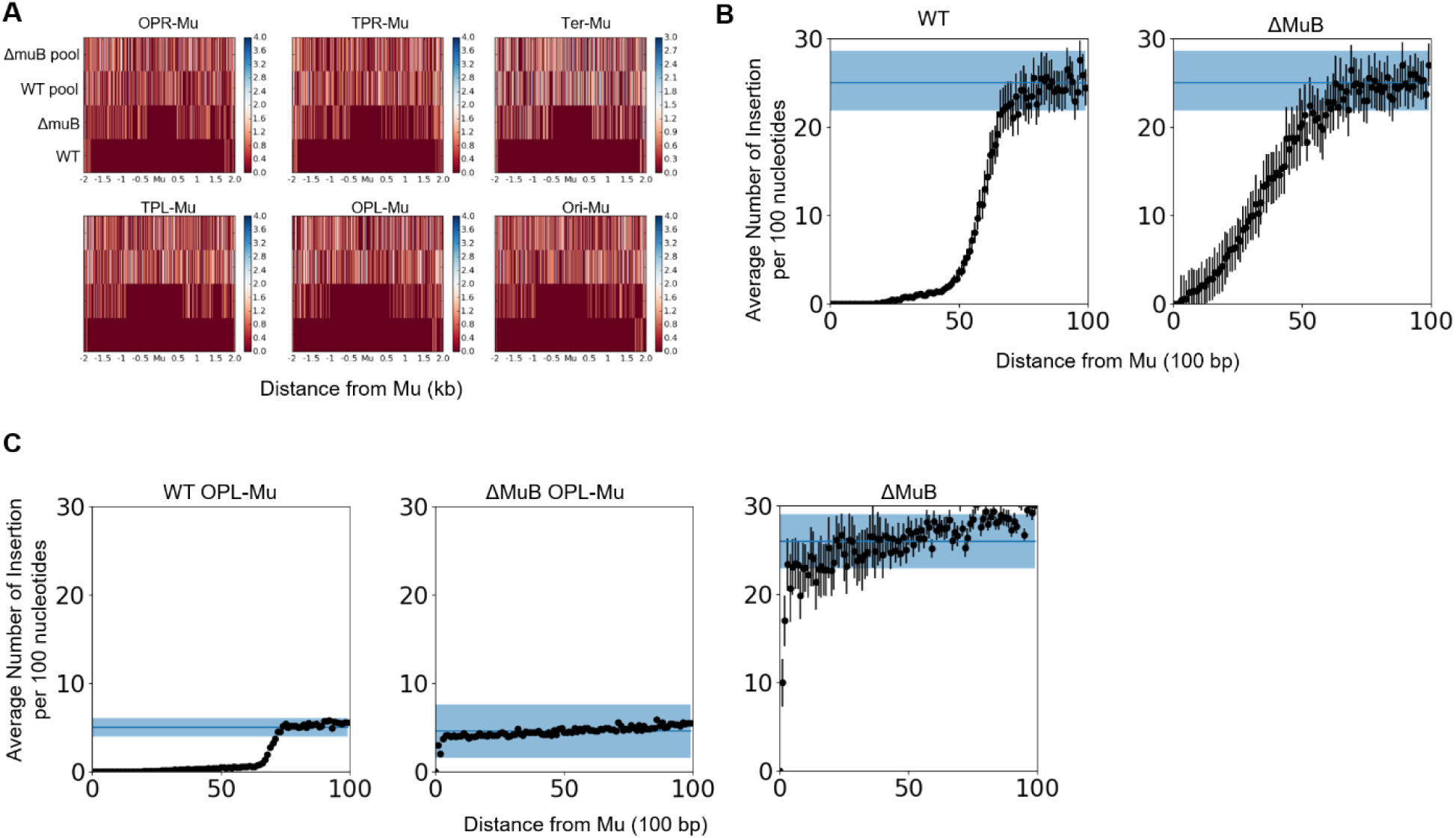
MuB is responsible for *cis*-immunity. The number of insertions near the initial starting location for each Mu prophage was tracked outside both the left and right ends of Mu during EST(**A** and **B**) and LST (**C**). **A**. The frequency of Mu insertions during EST for all six prophages under four different experimental steps are shown. Pooled experiments are frequency of insertions into that particular location from the other 5 prophages, and indicate that all these particular chromosomal locations are readily transposed into in the absence of Mu. **B)** The frequency of Mu insertions per 100 bp as a function of distance outside Mu. For bulk DNA the average number of insertions into a 100 bp region is nearly 25 insertions per 5 million reads and is indicated by the solid blue line. The distances reported are combined for both the left and right ends of Mu (see Fig. S4 for individual ends). The shaded blue area is the standard deviation for the number of insertions expected within 100 bp. **C**. The frequency of Mu insertions during LST. The number of Mu insertions outside both the left and right are reported as in **B** for WT OPL-Mu (left), ΔMuB OPL-Mu (middle), and all 6 ΔMuB prophages (right).

In the absence of MuB, we expected to see a maximal insertion frequency (consistent with that for bulk DNA) around 150 bp outside Mu ends, which is the *in vitro* persistence length of DNA [52]. Instead, EST ΔMuB prophages exhibited a more gradual increase in insertion frequency, starting between 500 to 600 bp and reaching bulk transposition efficiency around 7 kb from the ends (Fig. 4B and Fig. S4). This pattern was different in the LST samples, where *cis-*immunity remained intact for the WT OPL-Mu (Fig. 4C left), but was completely abrogated in ΔMuB OPL-Mu alone (compare Fig. 4C middle with left), and well as in all six ΔMuB prophages combined (Fig. 4C right). For the ΔMuB prophages, insertions started at 98 bp (at a distance smaller than the *in vitro* persistence length; [53]) and reached bulk efficiency between 2 and 6 kb. Why are the ΔMuB insertion patterns so different between the EST and LST samples? We think the lower transposition efficiency of ΔMuB prophages did not provide sufficient opportunity to sample nearby space during EST, whereas the increased ΔMuB copy number in the LST samples provided a greater opportunity to saturate the *cis* region. We conclude that MuB is indeed responsible for *cis*-immunity *in vivo*.

What is the importance of *cis*-immunity in the life of Mu? Avoiding insertion into regions flanking Mu ends would avoid destroying flanking Mu copies when packaging begins, since the DNA packaging machinery resects on average 100 bp of host DNA flanking the left end and 1.5 kb of DNA flanking the right end. Negating this concern, however, is the finding that Mu samples the *E. coli* genome extensively in a distance-independent manner (Fig. 2) [29]. A more likely possibility is that *cis*-immunity is an evolutionary remnant of MuB- and MuA-like functions in an ancestral transposon, where additional partner proteins directed transposition to specific sites. For example, Tn3 and Tn7 exhibit target immunity much further than Mu [22, 54, 55]. Tn7 has two proteins TnsB and TnsC that are thought to play roles similar to MuB and MuA respectively. Tn7 has two partner proteins, TnsD and TnsE, that promote different target choices. Han and Mizuuchi [25] discuss how the Mu *cis*-immunity system may have evolved from a Tn7-type target site search. Mu apparently discarded these partners during an evolutionary trajectory more suited to its viral lifestyle, acquiring features that unfettered its ability to choose.

### MuB is only partially responsible for Mu genome immunity

The *cis*-immunity phenomenon depends on MuB removal from DNA adjacent to and outside Mu ends. By contrast inside Mu, the MuB was observed to bind strongly during the lytic cycle, implicating a role for bound MuB in Mu genome immunity [27]. In the EST insertion data shown in Fig. 4A, there were no observable self-insertions (SI) in either WT or ΔMuB (the latter have 1.5x the depth of sequence reads compared to WT). SI was also not detected in the EST data for 35 WT prophages reported earlier [29]. To determine if this immunity is still intact at the end of the lytic cycle, we examined LST counts in the two prophage populations (Fig. 5). The WT OPL-Mu was still immune to SI (not shown), but the ΔMuB prophages, which have higher copy numbers in LST, now showed evidence of self-insertion. However, out of 90 Million Mu targeted reads from deep sequencing, 85 instances of SI were observed, spread across all 6 starting ΔMuB prophages. We conclude that, unlike *cis*-immunity which is completely abrograted in the absence of MuB (Fig. 4C), genome immunity is only slightly violated. Therefore, the bulk of genome immunity is determined by factors other than MuB.

**Figure 5.**
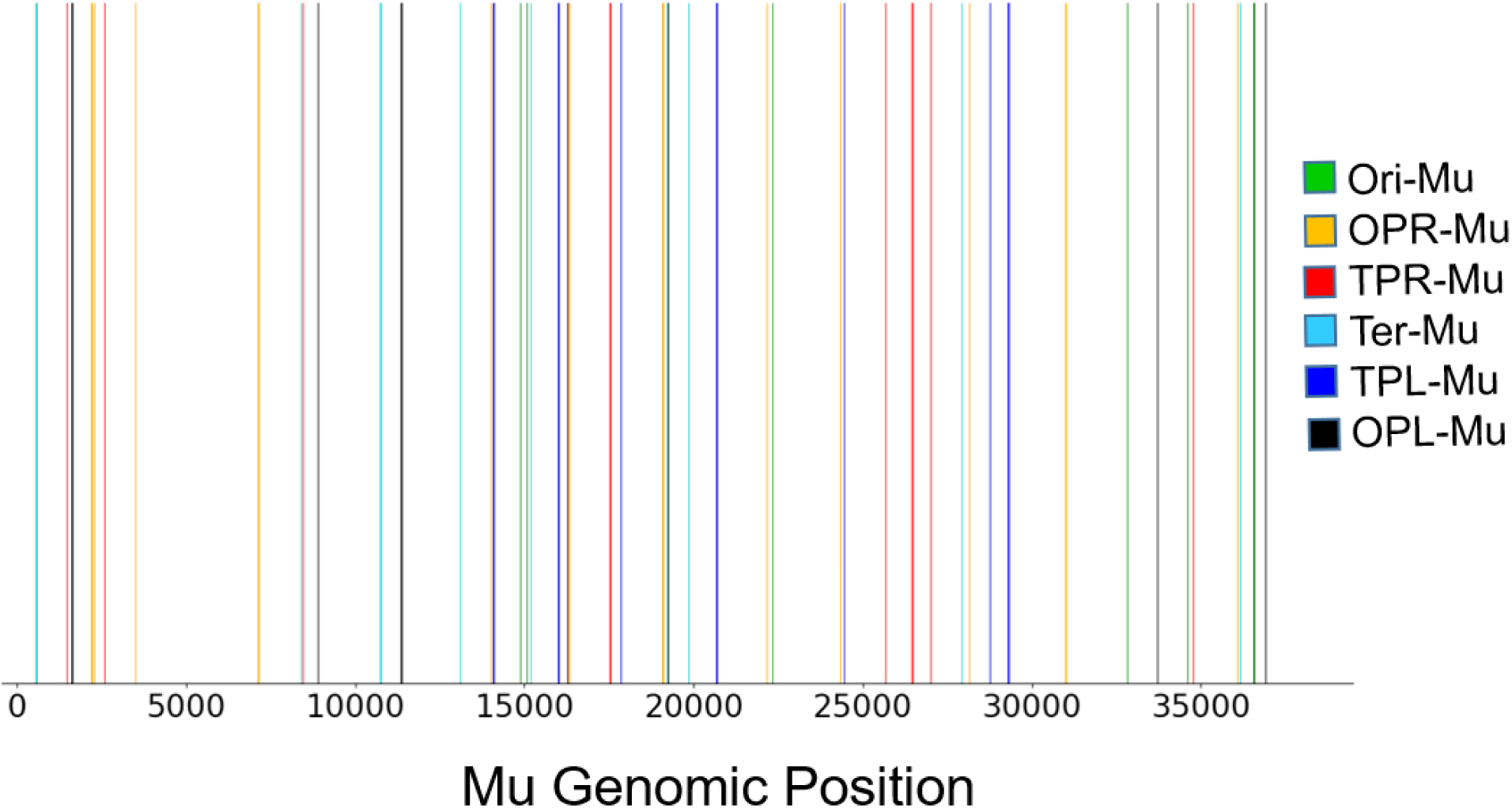
ΔMuB prophages exhibit very low levels of self-integration. Deep sequencing WT and MuB LST prophages were analyzed for novel Mu junctions that would indicate Mu self-integration. Out of ∼10 Million WT, no instances of self-integration were observed (data not shown). The 85 Mu self-integration sites observed in ΔMuB LST prophages are plotted along the Mu genomic position. Each insertion is color coded to correspond which prophage that specific insertion belongs to.

Mu ends (L and R) define a boundary separating two modes of MuB binding and immunity [27]. We had proposed that Mu genome immunity arises from a special structure that Mu adopts, aided by both specific Mu sequences and by general cellular NAPs. In the center of the genome is the strong gyrase-binding site (SGS), which is essential for Mu replication *in vivo* and is believed to function by influencing efficient synapsis of the Mu ends [56-58]. The SGS is thought to act by localizing the 37 kb Mu prophage DNA into a single loop of plectonemically supercoiled DNA upon binding of DNA gyrase to the site. We had proposed that an SGS-generated Mu loop, sealed off at the Mu ends by either the transpososome or NAPs, serves as a scaffold for nucleating MuB filaments in the Mu interior, providing a barrier to Mu integration. Evidence for a separate, stable prophage Mu domain, bounded by the proximal location of Mu L and R ends, was indeed obtained [28]. Formation/maintenance of the Mu domain was dependent on SGS, the Mu L end, MuB protein, and the *E. coli* NAPs IHF, Fis and HU. Of these components, SGS is essential for Mu transposition *in vivo* [59, 60], hence its contribution to Mu genome immunity cannot be assessed. To examine the contribution of the NAPs, we analyzed our published data where we had monitored Mu transposition in all NAP mutants of *E. coli* (these were collected during EST) [29]. We observed no instances of Mu self-transposition in any of the NAP deletions examined.

### Summary

MuB is critical for Mu’s ability to efficiently capture targets for transposition. We show in this study that besides enabling efficient targeting, MuB also makes refractory targets more facile, likely by displacing bound proteins. By weakening/altering boundary features that demarcates the Ter region, MuB allows Mu to access Ter more readily. Transposition patterns in the absence of MuB have allowed us to more accurately measure the Ter boundaries, revealing that this region is larger than previously estimated. Perhaps in a similar manner, MuB also provides access to targets engaged in transcription/translation. We have mapped the range of *cis*-immunity more accurately, and show that it persists well into the lytic cycle for WT prophages, but is abolished in ΔMuB strains. We show that Mu genome immunity also persists through the lytic cycle for WT prophages, and is only rarely infringed upon in ΔMuB prophages, showing conclusively the distinction between these two forms of immunity. There is clearly more to be learned about what enables genome immunity.

## Materials and Methods

### Strain Information and Growth Conditions

All experimental strains are derivatives of MG1655 and listed in the strains table. Prophage gene deletions were introduced into specific prophages using P1 transduction and kanamycin resistance selection. Cells were propagated by shaking at 30 °C in M9-Cas minimal media (0.2% casamino acids, 0.2% glucose, 100 ug/mL thiamine) and appropriate antibiotics for selection.

### Transposition

Prophage transposition was induced by temperature shifting to 42 °C for the appropriate time before harvesting genomic DNA. Early stage transposition (EST) experiments were accomplished by a 15 minute temperature shift to capture one transposition event in WT cells as determined in a previous study [29]. Late stage transposition (LST) experiments were done by a temperature shift for 2 hours. At the end of this time, cell lysis had occurred for WT prophages but not for ΔMuB prophages. Lysogen genomic DNA was purified using a commercially available gDNA purification kit (Wizard, Promega). gDNA samples were stored at -20 °C in a 10 mM Tris pH 8.0, 1 mM EDTA buffer until ready for target enrichment.

### Target Enrichment

Primer y-link1 has a hand mixed random 6 nucleotide barcode to identify PCR duplicates in sequencing. Y-link adapters were annealed by mixing equivalent amounts of primers y-link1 and y-link2 at room temperature and heating to 95 °C then cooled down to 4 °C using a temperature ramp of 1 °C per second. Genomic DNA was digested with the frequent cutter HinPI (NEB) and then ligated with the y-link adapter using a quick ligase kit (NEB). The ligation product was purified using magnetic beads (Axygen). Mu insertion targets were enriched, by PCR amplification of the ligation product using y-link_primer and Mu_L31, an initial melting temp of 95 °C for 1 min and 8 cycles of 95 °C for 20 s, 68 °C for 20 s, 72 °C for 1 min. A final extension of 72 °C was added for 5 minutes. The PCR product was purified using magnetic beads (Axygen) and frozen at -20 °C until ready for sequencing.

### Genomic Sequencing

Target enriched samples were submitted to the Genomic Sequencing and Analysis Facility (GSAF) at UT Austin for sequencing. Libraries were prepped by GSAF using the facility’s low-cost high throughput method. Sequencing was done on an Illumina NextSeq 500 platform using 2×150 paired ends targeting 10 to 15 million reads. All sequencing data discussed in this work is available at https://www.ncbi.nlm.nih.gov/sra/PRJNA597349.

### Identifying Mu Insertion Locations

Mu transposition targets were identified using lab software entitled Mu Analysis of Positions from Sequencing (MAPS) as described earlier [29]. MAPS has been modified since initial publication to provide nucleotide precision for target enriched samples and provide self-insertion information. In short, MAPS now identifies Mu-host junctions by identifying a 12-mer sequence unique to the y-link adapter used in target enrichment. The current version of MAPS is available for download at https://github.com/dmwalker/MuSeq.

## Declarations

### Availability of data and material

All strains generated in this study are available without restriction. The sequencing data presented in this paper can be accessed on the SRA database under the project number PRJNA597349. Software used to analyze the sequencing data can be accessed by github (DOI 10.5281/zenodo.3762807).

### Competing interests

None

### Funding

National Institutes of Health (GM118085).

### Author contributions

DMW performed the experiments, DMW and RMH analyzed the data and wrote the manuscript.

## Acknowledgements

We thank Brady Wilkins and Mark Faulkner for strain construction and genomic DNA isolation, and Peter Freddolino for helpful comments on data analysis. Computer time was provided via the Texas Advanced Computing Center (TACC).

## Supplementary Figures

**Figure S1.**
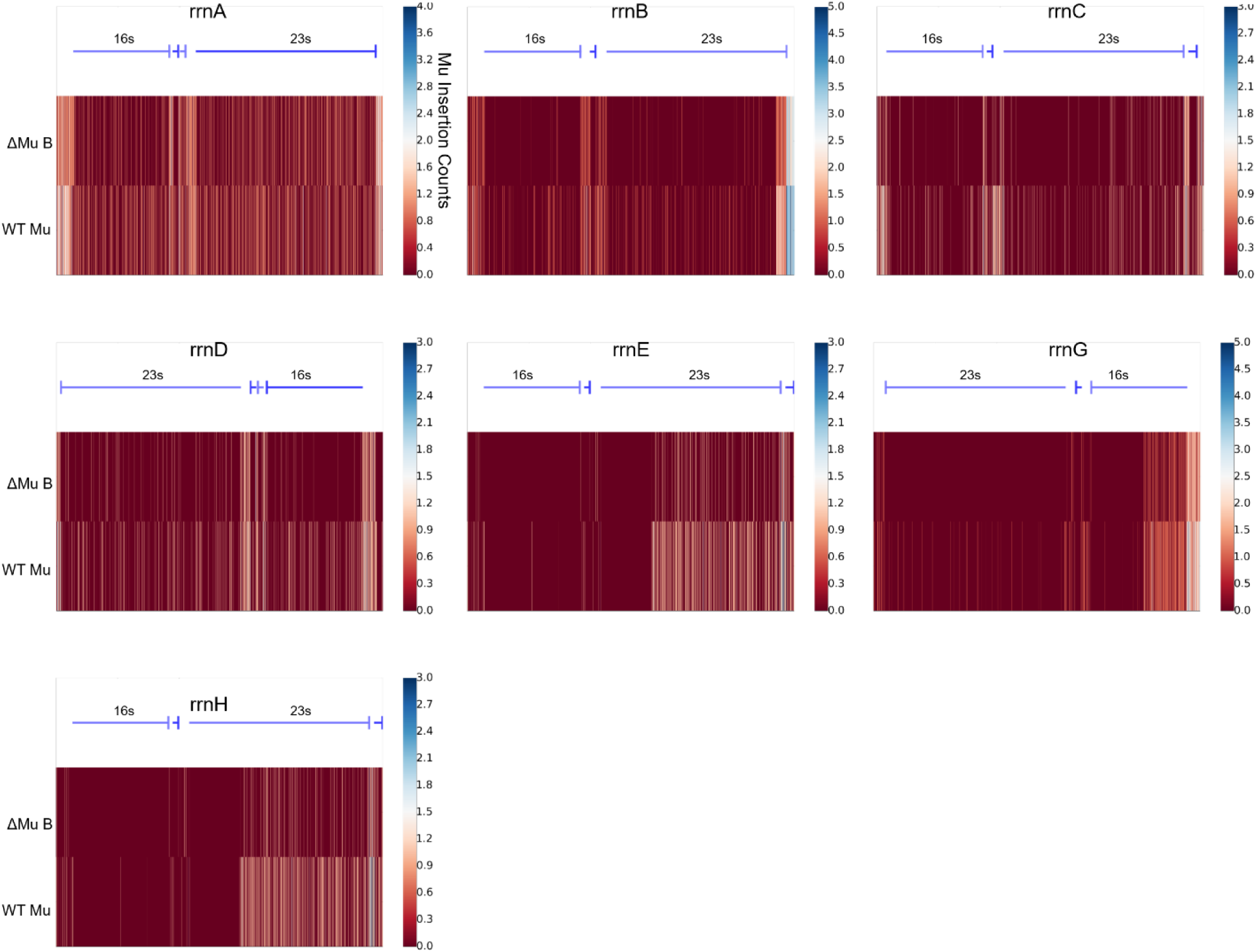
Mu transposition outlines several features of the rRNA operons. EST prophages were pooled to analyze the frequency of Mu insertions into the entire *rrn* operon, for all 7 operons, for both WT (bottom rows) and ΔMuB (top rows) prophages. Insertion maps start at the TSS of the operon and continue for 5.3 kb. Operon maps are provided as a schematic on top, showing the leading 16s RNA-encoding segment, followed by coding sequence (CDS) of an intervening tRNA, and finally the 23s RNA-encoding segment. Each CDS in the operon is marked by a blue line that terminates in a flat head. The *rrnD* and *rrnG* operons are located on the (-) strand of DNA, while the remaining 5 are on the (+) strand. ΔMuB patterns follow similar trends to the WT prophage, but with reduced efficacy to insert anywhere within the rRNA operon.

**Figure S2.**
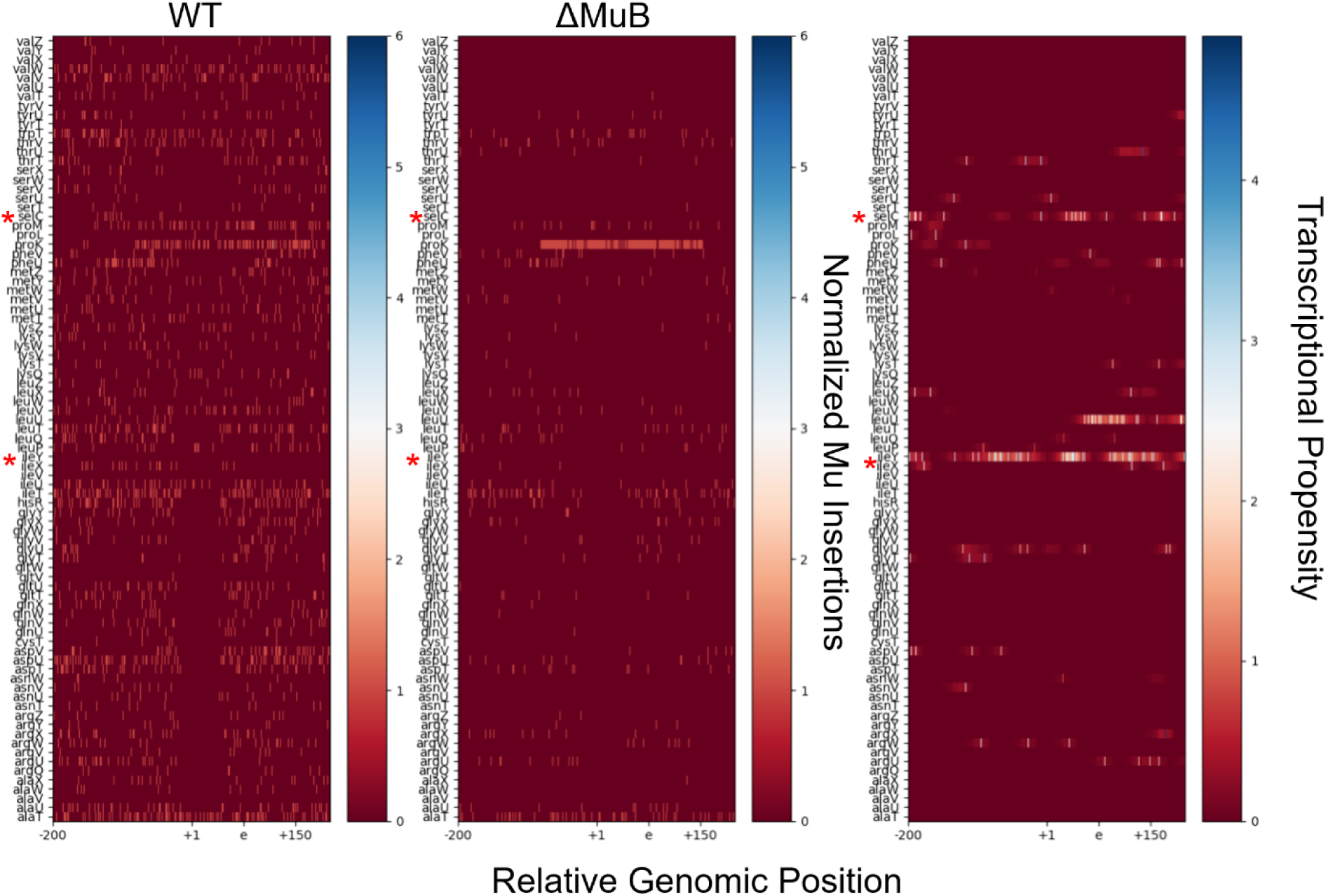
Mu does not transpose easily into tRNA coding regions. The x-axis provides the relative genomic position with respect to the tRNA labeled on the y-axis, and covers a 400 nucleotide span. The +1 position indicates the first nucleotide in the matured tRNA sequence. -200 nt from the mature tRNA +1 position and would contain the preprocessed 5’ leader. The e position is +75 nt from the TSS and is the typical size of mature tRNA. The +150 region is 150 nucleotides from the TSS. For each of the 86 tRNA genes, the number of Mu insertions in and around the gene are tabulated for both the WT and ΔMuB prophages during EST. The transcriptional propensity is nucleotide level resolution of the degree of transcription for that particular nucleotide [2]. A higher number means higher degree of transcription.

**Figure S3.**
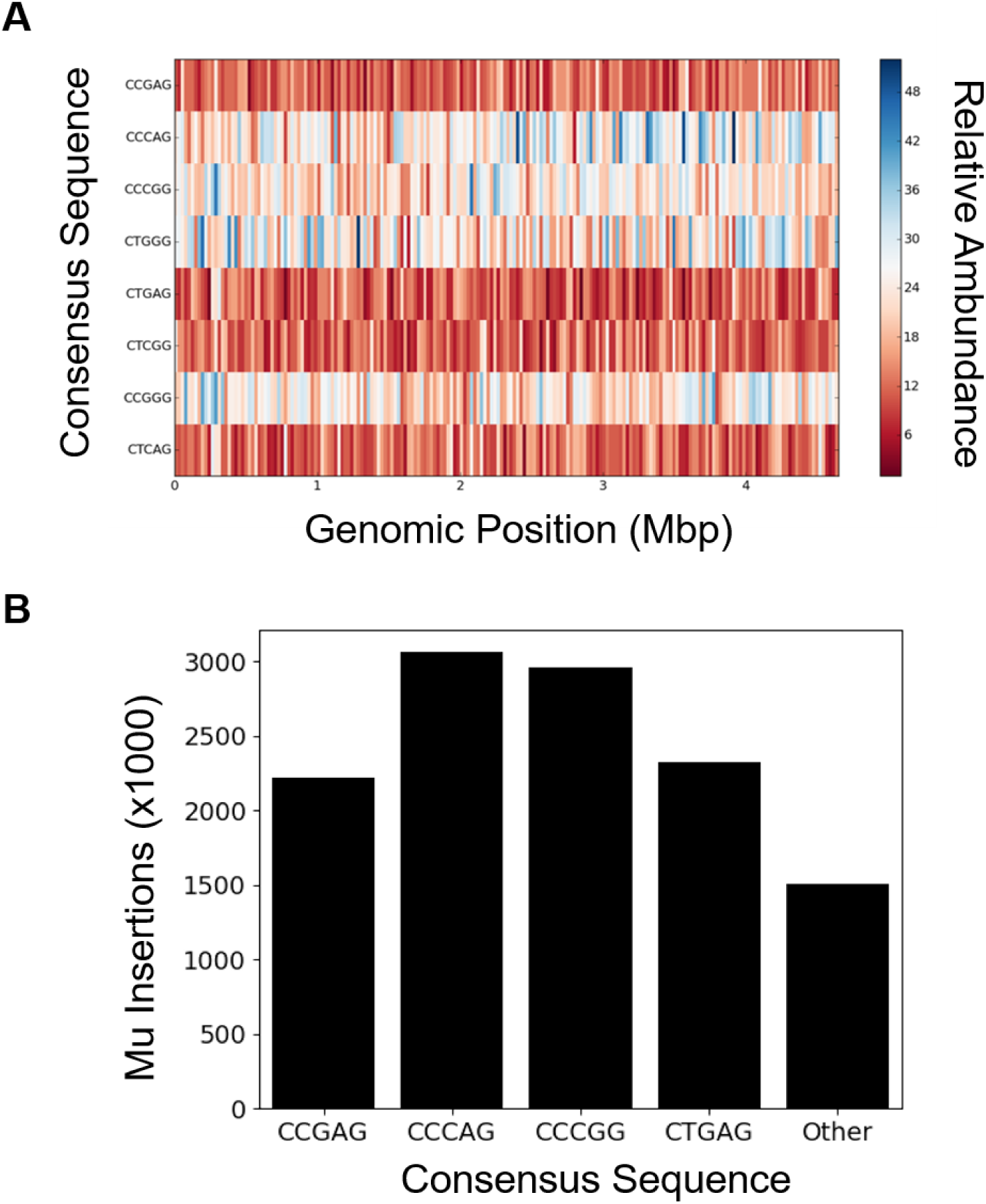
Frequency of consensus target sequences for WT Mu across the *E. coli* genome. **A**. The genome for MG1655 from genbank (genid: 545778205) was partitioned into 200 equally sized bins, and the number of times the 5’-CYSRG-3’ sequence and it’s reverse compliment appeared on the + strand in each bin was tabulated. **B**. The number of Mu insertions for each consensus sequence was calculated. The number of insertions reported is for the consensus sequence written and the corresponding reverse compliment. There are 1024 possible pentamers for Mu to insert, and the sequence identifier ‘other’ accounts for the 1016 sequences not covered by the ‘CYSRG’ consensus sequences and their reverse compliment.

**Figure S4.**
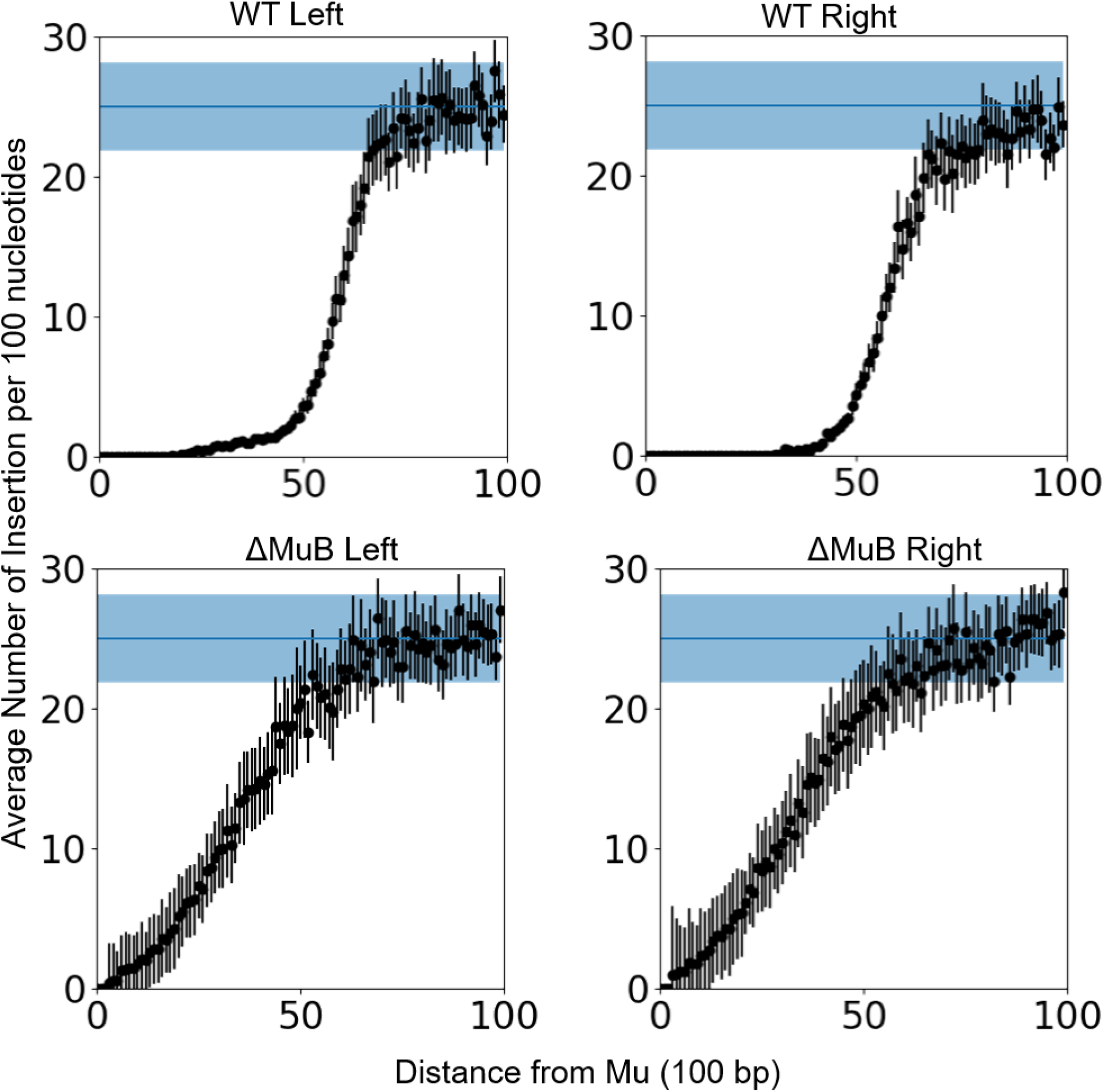
Insertion patterns outside Mu ends are nearly symmetrical for the left and right ends of Mu. The frequency of Mu insertions per 100 bp as a function of distance from Mu is plotted individually for each end, the combined data shown in Fig. 4. For WT Mu, the first Mu insertion on the left side occurred at 1.6 kb from the left end, while ΔMuB insertions started at 529 bp. The right end insertions of WT Mu started around 3.1 kb and at 544 bp for ΔMuB prophages. Wild type *cis*-immunity shows a sharp decline around 5Kb for both the right and left ends of Mu, while the ΔMuB prophages show a steady increase in insertions away from initial prophage.

## Key Resources Table

| REAGENT or RESOURCE | SOURCE | IDENTIFIER |
| --- | --- | --- |
| Chemicals, Peptides, and Recombinant Proteins |  |  |
| HinP1 | New England Biolabs | R0124S |
| Phusion Polymerase | New England Biolabs | M0530S |
| Critical Commercial Assays |  |  |
| Wizard Genomic DNA Purification Kit | Promega | A1120 |
| Quick Ligation Kit | New England Biolabs | M2200L |
| Qiaquick PCR Cleanup Kit | Qiagen | 28106 |
| Axygen™ AxyPrep Mag™ PCR Clean-up Kits | Thermo Scientific | 14-223-227 |
| Deposited Data |  |  |
| Genomic Sequencing Data | This Study | https://www.ncbi.nlm.nih.gov/sra/PRJNA597349 |
| Organisms/Strains |  |  |
| MP1999 *Mucts B::kan*. Mu insertion between nts 365204-3652051 | (Saha et al. 2013) | ZL530 |
| MG1655 with *Mucts:cat* at nt 4203381 (Ori-Mu) | (Walker et al. 2020) | DMW11 |
| MG1655 with *Mucts:cat* at nt 263131 (OPR-Mu) | (Walker et al. 2020) | DMW15 |
| MG1655 with *Mucts:cat* at nt 3339450 (OPL-Mu) | (Walker et al. 2020) | DMW22 |
| MG1655 with *Mucts:cat* at nt 2555144 (TPL-Mu) | (Walker et al. 2020) | DMW24 |
| MG1655 with *Mucts:cat* at nt 792226 (TPR-Mu) | (Walker et al. 2020) | DMW33 |
| MG1655 with *Mucts:cat* at nt 1657887 (Ter-Mu) | (Walker et al. 2020) | DMW57 |
| DMW57, *Mucts:cat B::kan* | This study, DMW57, ZL530 | DMW300 |
| DMW33, *Mucts:cat B::kan* | This study, DMW33, ZL530 | DMW301 |
| DMW24, *Mucts:cat B::kan* | This study, DMW24, ZL530 | DMW302 |
| DMW22, *Mucts:cat B::kan* | This study, DMW22, ZL530 | DMW303 |
| DMW15, *Mucts:cat B::kan* | This study, DMW15, ZL530 | DMW304 |
| DMW11, *Mucts:cat B::kan* | This study, DMW11, ZL530 | DMW305 |

| Software and Algorithms |  |  |
| --- | --- | --- |
| MAPS (Python) | This Study | https://github.com/dmwalker/MuSeq |
| BWA-MEM | Li H., 2013 | https://sourceforge.net/projects/bio-bwa/ |

## Notes

### Competing Interest Statement

The authors have declared no competing interest.

